# Cytotoxicity of Pelargonic Acid and Its Commercial Formulation Roundup NL (Glyphosate-Free Roundup)

**DOI:** 10.64898/2026.07.07.736979

**Authors:** Scarlett Ferguson, Robin Mesnage, Michael N Antoniou

## Abstract

Evidence of negative health and environmental effects of glyphosate-based herbicides (GBHs) has led to marketing of glyphosate-free formulations. A frequent glyphosate replacement is pelargonic acid, which is rapidly degraded, leading to claims of greater safety and less environmentally damaging than GBHs. However, toxicity of commercial pelargonic acid formulations containing several co-formulants have not been determined. Using Roundup NL, a representative pelargonic acid-based herbicide, we undertook tissue culture cell assays measuring viability, plasma membrane integrity, DNA damage, and activation of stress-response pathways. In human hepatoma HepG2 cells, Roundup NL was more cytotoxic than pelargonic acid, and more toxic than the GBH Roundup ProBio and glyphosate as shown by reduced viability underpinned by plasma membrane damage. Pelargonic acid and Roundup NL did not induce oxidative stress. However, comet assays revealed that pelargonic acid but not Roundup NL caused a modest but significant increase in DNA damage at sub-cytotoxic concentrations. The murine embryonic stem cell-based ToxTracker system confirmed Roundup NL as not directly genotoxic but triggered oxidative stress and protein damage (ER stress, impaired proteostasis) indicating cell and assay dependency of oxidative stress pathway activation. Our results suggest that exposure to pelargonic acid-based herbicides constitutes a health hazard and that co-formulants present in Roundup NL contribute substantially to its overall toxicity.

## 1. Introduction

Glyphosate is among the world’s most widely used herbicides, owing to its broad-spectrum efficacy and extensive adoption in agriculture, especially with glyphosate-tolerant genetically modified crops (Mesnage and Antoniou, 2021). Since its introduction in the 1970s, in excess of several billion kilograms of glyphosate have been applied worldwide with annual use currently being estimated at approximately 749 million kg (Benbrook, 2016, Maggi et al., 2020). However, the pervasive use of glyphosate-based herbicides (GBHs) has become highly controversial. On the one hand, regulatory agencies in many jurisdictions such as the United States (US EPA, 2026), European Union (EU, 2023) and Canada (Health Canada, 2017) have repeatedly concluded that glyphosate poses no significant human health risk when used as directed. On the other hand, in 2015 the International Agency for Research on Cancer classified glyphosate herbicides as “probably carcinogenic to humans”, citing epidemiological and toxicological studies (Guyton et al., 2015). An increasing body of evidence indicates that GBH exposure can not only be carcinogenic (Weisenburger, 2021, Weisenburger, 2026, Panzacchi et al., 2025), but can also be a risk factor for the development of fatty liver disease (Riechelmann-Casarin et al., 2025), neurological defects (Costas-Ferreira et al., 2022) and potential endocrine-disrupting effects (Muñoz et al., 2021; Serra et al., 2021). The EU acceptable daily intake of glyphosate (0.5mg/kg body weight/day), which is half that set by the Joint Meeting on Pesticide Residues/World Health Organisation (JMPR, 2016), and adopted by most other countries, still resulted in carcinogenic outcomes in a regulatory compliant investigation in rats (Panzacchi et al., 2025), questioning current safety limits of glyphosate exposure. In addition, the large, widespread use of GBHs can lead to negative ecological impacts (Klátyik et al., 2023, Klátyik et al., 2024, Klátyik et al., 2025). This dichotomy between industry/regulatory assessments and independent evaluations has fuelled an intense public debate over GBH safety. In response to mounting public pressure and legal liabilities, GBH manufacturers have begun seeking alternatives to glyphosate.

One prominent replacement of glyphosate as an active herbicidal ingredient is pelargonic acid (nonanoic acid). Pelargonic acid is a naturally occurring fatty acid present in many plants and common foodstuffs (e.g. apples, grapes, cheese, milk, rice, beans, oranges, potatoes) and therefore part of the human diet (US EPA, 1998). Pelargonic acid has a very different mode of action from glyphosate being a fast-acting contact herbicide that disrupts cell membranes and cuticular waxes resulting in rapid plant death (Preisler et al., 2026). Because pelargonic acid is naturally found in plants and readily biodegradable, regulators such as the US Environmental Protection Agency (EPA) categorise it as a biopesticide, meaning it is considered to pose lower risk than conventional synthetic herbicides (US EPA, 2003). This status has opened the door for marketing glyphosate-free herbicides as “natural” or “safe” – for example, pelargonic acid-based products are often advertised as pet-safe and environmentally benign. The European Food Safety Authority (EFSA) recently concluded that pelargonic acid is unlikely to be genotoxic and did not identify serious health hazards at expected exposure levels (EFSA, 2025). Given this ostensibly benign profile, pelargonic acid-based herbicides are widely assumed to be safer alternatives to GBHs.

Despite the presumed safety of pelargonic acid itself, there is a notable lack of toxicity evaluation for the full commercial formulations of glyphosate-free products containing this compound. Pesticides are almost never applied as pure active ingredients; they are formulated with various surfactants, solvents, and adjuvants to enhance efficacy (Mesnage and Antoniou, 2018). In this regard commercial formulations of pelargonic acid-based herbicides are no exception and contain surfactants and emulsifiers as co-formulants to enhance solubility, wettability and uniform leaf adhesion (Preisler et al., 2026). These co-formulants are typically labelled “inert” from a regulatory standpoint, but a large and increasing body of evidence suggests many are not toxicologically inert. In the case of GBHs, studies have shown that co-formulants (e.g. ethoxylated tallowamine surfactants) can markedly increase toxicity to non-target cells (Mesnage et al., 2013; Mesnage et al, 2019). However, for the new glyphosate-free herbicides, similar scrutiny has lagged. Many of these products were fast-tracked to market on the assumption that replacing glyphosate with a “natural” acid ensures safety, with little requirement to disclose or test the full commercial mixture’s toxicity.

In light of the above, we set out to investigate cellular cytotoxic and stress-inducing effects of a pelargonic acid-based, glyphosate-free commercial herbicide formulation, Roundup NL, and compared this to pelargonic acid alone, glyphosate and a conventional GBH. We employed a battery of contemporary *in vitro* tissue culture assays encompassing cell viability, membrane integrity, DNA damage, and activation of stress-response pathways. Our findings provide important new evidence regarding the safety of a pelargonic acid-based glyphosate-free herbicide that is marketed as a safer and environmentally friendly product.

## 2. Methods

### 2.1 Herbicide Materials

Glyphosate (N-(phosphonomethyl)glycine and pelargonic acid were of analytical grade and obtained from Merck Life Sciences (Watford, UK). The commercial formulations used were Roundup ProBio (Monsanto UK Ltd., Cambridge, UK; 360 g/L glyphosate) and Roundup NL (Evergreen Garden Centre, Surrey, UK, 43.1g/L pelargonic acid). Stock solutions of the test substances were prepared in dimethyl sulphoxide (DMSO). Dilutions of the stock solutions administered to cells resulted in concentrations of DMSO below 0.05%. Serial dilutions of the active ingredients and formulations were prepared in appropriate tissue culture cell media.

### 2.1 HepG2 Cell Culture

The human hepatoma HepG2 cell line was obtained from the European Collection of Authenticated Cell Cultures (ECACC) and was used between passages 53 and 65. HepG2 cells were grown in high glucose, pyruvate Dulbecco’s Modified Eagle Medium (DMEM) (Thermo Fisher Scientific, Loughborough, UK) supplemented with 10% foetal bovine serum (South American origin, Thermo Fisher Scientific), 2 mM L-glutamine (Thermo Fisher Scientific) and 100 units penicillin/mL and 100 µg/mL streptomycin (Thermo Fisher Scientific). Cells were cultured in 75 cm^2^ flasks (Corning, Tewksbury, MA, USA) under standard conditions of 37°C and 5% CO_2_ air atmosphere. Depending on density, HepG2 cells were passaged every 3-4 days.

### 2.2 HepG2 Cell MTT Assay

Cell viability was assessed using a 3-(4, 5-dimethylthiazol-2-yl)-2, 5-diphenyltetrazolium bromide (MTT) assay. The MTT assay is a colorimetric assay based on the intracellular reduction of yellow MTT to purple formazan granules, testing for the activity of mitochondrial succinate. HepG2 cells were seeded at 50,000 cells/well in 100 µL of the medium in clear-walled 96 well plates (Corning, Thermo Fisher Scientific). Following a 24-hour incubation, cells were treated with the test substances diluted according to the desired concentrations in a tissue culture maintenance medium. After a further 24-hour incubation, an MTT assay was performed to assess cell proliferation and thus cytotoxicity according to the manufacturer’s instructions. Cells were incubated for 2 hours in MTT solution at 1 mg/mL in PBS. The resulting formazan precipitate was then dissolved by the addition of 100 µL DMSO and quantified spectrophotometrically at 560 nm using a GloMax Multi Microplate Multimode Reader (Promega UK Ltd, Chilworth, Southampton, UK). Cell viability was expressed as a percentage relative to the negative control, untreated cell samples consisting of cell culture medium only.

### 2.3 HepG2 ToxiLight Assay

Cytotoxicity was assessed using the ToxiLight bioluminescent assay (Lonza, Slough, Berkshire, UK), which measures adenylate kinase (AK) release from damaged cells as a marker of membrane disruption and necrosis. HepG2 cells were seeded in black-walled 96-well plates at 50,000 cells/well and incubated for 24 hours. Cells were then treated with test substances at LC50, LC50/2, and (LC50/2)/2 concentrations, or the three highest MTT assay concentrations if LC50 was unavailable.

After treatment, 50 µL of AK reagent was added per well. Luminescence, proportional to AK activity and thus cell damage, was measured after 5 minutes using a GloMax plate reader.

Results were expressed as relative light units (RLU), background-subtracted, and normalized to untreated controls. Triton X-100 (0.05%) served as a positive control for membrane rupture.

### 2.4 HepG2 ROS-Glo™ H_2_O_2_ Assay

Oxidative stress was measured using the ROS-Glo™ H_2_O_2_ bioluminescent assay (Promega), which quantifies hydrogen peroxide (H_2_O_2_) production in cell culture as a marker of reactive oxygen species (ROS). HepG2 cells were seeded at 10,000 cells/well in 96-well white-walled plates and incubated for 24 hours. Cells were then treated with test compounds or 50 µM menadione (positive control); untreated cells served as the negative control.

Immediately after treatment, 20 µL of H_2_O_2_ substrate was added. After 6 hours, 100 µL of ROS-Glo detection reagent was added per well following the manufacturer’s instructions. After a 20-minute incubation in the dark at room temperature, luminescence was measured using a GloMax plate reader. Results were reported as relative light units (RLU), reflecting H_2_O_2_ levels.

### 2.5 HepG2 Comet DNA Damage Assay

DNA damage was evaluated in HepG2 cells using the alkaline comet assay (Singh et al., 1988) employing the CometChip system of Bio-Techne Ltd (Abingdon, Oxfordshire, UK), which detects single- and double-strand DNA breaks. Cells were seeded at 200,000 cells/well in 12-well plates, incubated for 24 hours, and treated in duplicate with test compounds for 24 hours at LC10 concentrations, or the highest non-cytotoxic dose from the MTT assay when LC10 was unavailable. Hydrogen peroxide (50 µM, 30 min) served as a positive control, and untreated cells as the negative control.

Following treatment, cells were detached using 0.5% trypsin-EDTA, embedded in low melting point agarose on CometSlides™, and subjected to overnight lysis at 4□°C. Slides were then incubated in alkaline unwinding solution (pH >13) for 20 minutes and electrophoresed for 30 minutes at 21 V in alkaline electrophoresis buffer. After washing and ethanol fixation, slides were stained with SYBR® Gold and dried.

Fluorescence microscopy was used to image and quantify at least 100 comets per treatment. DNA damage was assessed as the percentage of DNA in the tail using CometAssay Analysis Software (Bio-Techne). Comet assays for each chemical were repeated twice in different weeks, with at least 200 comets per chemical scored (100 per slide).

### 2.6 ToxTracker® Assay

The ToxTracker^®^ assay (Hendriks et al., 2016) was conducted under contract with Toxys BV (Leiden, Netherlands) to evaluate the genotoxic and stress-inducing potential of Roundup NL as previously described (Mesnage et al., 2021; Mesnage et al., 2022a; Mesnage et al., 2022b; Ferguson et al., 2022). This stem cell-based reporter assay system uses six murine embryonic stem (mES) cell lines, each expressing a GFP reporter responsive to specific cellular stress pathways, including DNA damage (Bscl2, Rtkn), p53 activation (Btg2), oxidative stress (Srxn1, Blvrb), and protein damage/ER stress (Ddit3).

Cells were seeded in gelatin-coated 96-well plates (50,000 cells/well) and exposed for 24 hours to five concentrations of each compound in a 2-fold dilution series, both without and with metabolic activation using S9 liver extract from aroclor1254-induced rats (0.25% S9 mix). GFP expression was measured by flow cytometry, and reporter induction was calculated relative to solvent controls. Cytotoxicity was assessed via cell viability and reported as the percentage of intact, viable cells compared to controls. Positive controls included cisplatin (DNA damage), diethyl maleate (oxidative stress), tunicamycin (protein stress), and aflatoxin B1 (S9 activation control). Final solvent concentrations did not exceed 1% DMSO or 10% water.

#### 2.7 Statistical analysis

Experiments were repeated a minimum of 3 times in different weeks (n = 9). Statistical differences were calculated using a Sidak multiple comparison test, which followed a one-way ANOVA comparing treatment groups to the negative control using the GraphPad Prism 9 software package (version 9.4.1). ANOVA was first chosen to check for overall group differences (negative control, positive control, active ingredient, formulation). Then, Šidák tests for pairwise comparisons was undertaken, if assumptions of ANOVA are violated (i.e. if differences amongst groups exist).

## 3. Results

We began our investigation by assessing the general cytotoxic potential of pelargonic acid and its representative commercial formulation Roundup NL on human hepatoma HepG2 cells. We chose the HepG2 cell line for these experiments as it is widely used in toxicology studies as an *in vitro* system to model systemic exposure effects and xenobiotic metabolism capacity.

Cell viability assessed by the MTT assay revealed that Roundup NL was more cytotoxic than pelargonic acid alone, with LC50 values of 320 µg/mL and 514 µg/mL, respectively (Figure 1). Roundup ProBio exhibited an intermediate cytotoxicity (LC50 = 440 µg/mL), while glyphosate did not reduce cell viability up to 500 µg/mL. Thus, Roundup NL was over 1.5 times more cytotoxic than pelargonic acid and more toxic than the glyphosate-based formulation it aims to replace. The establishment of the LC50 values for the various test substances guided the subsequent assays looking at other toxicity endpoints in HepG2 cells (Table 1).

**Table 1.** LC50, LC50/2 and (LC50/2)/2 concentrations of pelargonic acid, Roundup NL, glyphosate and Roundup ProBio used in ToxiLight membrane damage and oxidative stress assays.

| Substance | LC50, LC50/2 and (LC50/2)/2 Concentrations (µg/mL) |
| --- | --- |
| Pelargonic acid | 514, 257, 128.5 |
| Roundup NL | 320, 160, 80 |
| Glyphosate | 500, 250, 100 |
| Roundup ProBio | 440, 220, 110 |

**Figure 1.**
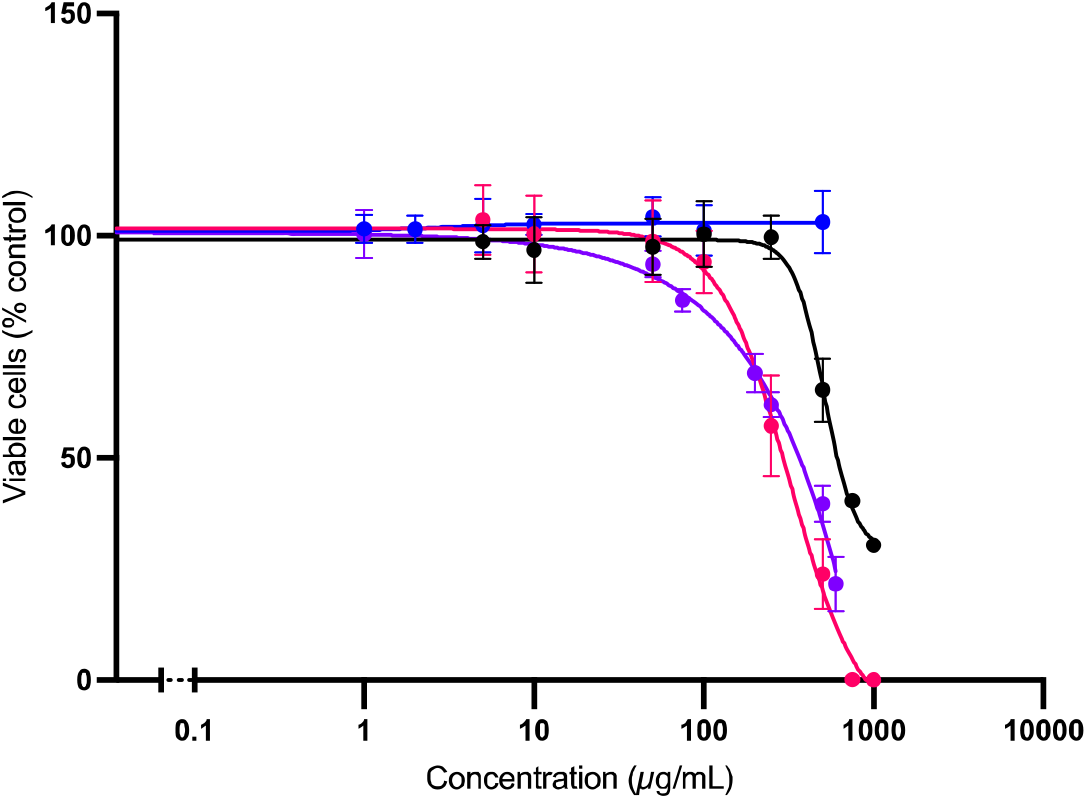
Differential cytotoxicity of pelargonic acid and Roundup NL in HepG2 cells. HepG2 cell proliferation and viability were assessed by MTT assay following exposure to pelargonic acid, Roundup NL, glyphosate and Roundup ProBio. Concentrations in µg/mL are dilutions of pelargonic acid (black), Roundup NL (pink), glyphosate (blue) and Roundup ProBio (purple). Cell viability was expressed as a percentage relative to the negative control, untreated cell samples. Roundup NL caused a greater decrease in cell viability compared to pelargonic acid alone. Pelargonic acid caused a greater reduction of viable cells compared to glyphosate. Roundup NL caused a greater reduction of viable cells than Roundup ProBio. The assay was performed in triplicate and data is expressed as mean ± SD (standard deviation) of 3 independent replicates.

Assessment of effects on HepG2 cell membrane integrity using the ToxiLight assay was undertaken at the LC50, LC50/2 and (LC50/2)/2 concentrations of pelargonic acid and Roundup NL as well as glyphosate and Roundup ProBio GBH (Table 1).

HepG2 cell membrane disruption as measured by adenylate kinase release showed that pelargonic acid and Roundup NL significantly increased this outcome by approximately 100% at their respective LC50 values (Figure 2). In contrast, glyphosate and Roundup ProBio did not induce significant membrane damage at any concentration tested. Notably, Roundup NL triggered more membrane disruption than Roundup ProBio, despite containing a lower concentration of the active ingredient, underscoring the enhanced cytotoxic potential of the Roundup NL formulation.

**Figure 2.**
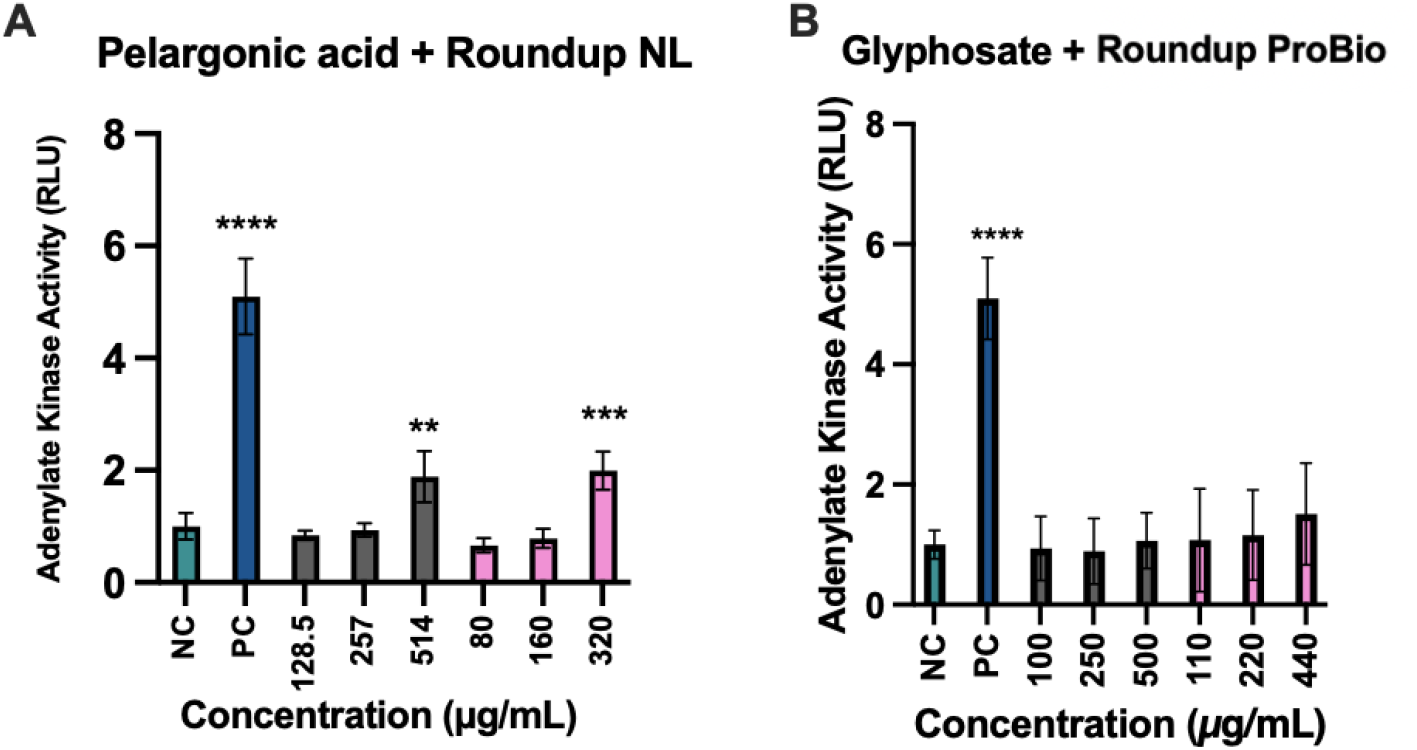
Measurement of membrane disruption cytotoxicity by pelargonic acid and glyphosate-free Roundup NL treatment in HepG2 cells. (**A**) Untreated negative control cultures (NC; green bars), positive control cultures (PC; blue bars) treated with 0.05% Triton X-100 were compared to pelargonic acid (grey bars) and glyphosate-free Roundup NL (pink bars) exposed cells at LC50, LC50/2 or (LC50/2)/2 at 24 h after exposure. Following treatment, cultures were assessed for plasma membrane damage and necrosis using the Toxilight assay system, which measures the release of adenylate kinase from damaged cells. The assay was performed in triplicate and data is expressed as mean ± SD (standard deviation) of at least 3 independent replicates. Adenylate kinase activity is expressed in relative units (RLU). Pelargonic acid and Roundup NL induce a significant increase in cell necrosis at their LC50 concentrations of 514μg/mL and 320μg/mL respectively. (**B**) Neither glyphosate nor Roundup ProBio induced and increase in cell necrosis at any concentration tested. Standard deviation (SD) is shown in all instances (n = 9). ** p <0.01, ***p <0.001, ****p <0.0001 in a Šidák test which followed a one-way ANOVA.

Measurement of oxidative stress in HepG2 cells via H_2_O_2_ production demonstrated that neither pelargonic acid, Roundup NL, nor Roundup ProBio induced significant ROS generation (Figure 3). Contrastingly, glyphosate elicited a concentration-dependent increase (50-100%) in H_2_O_2_ production at the LC50 and LC50/2 concentrations, indicating its greater potential to trigger oxidative stress compared to pelargonic acid and its formulation.

**Figure 3.**
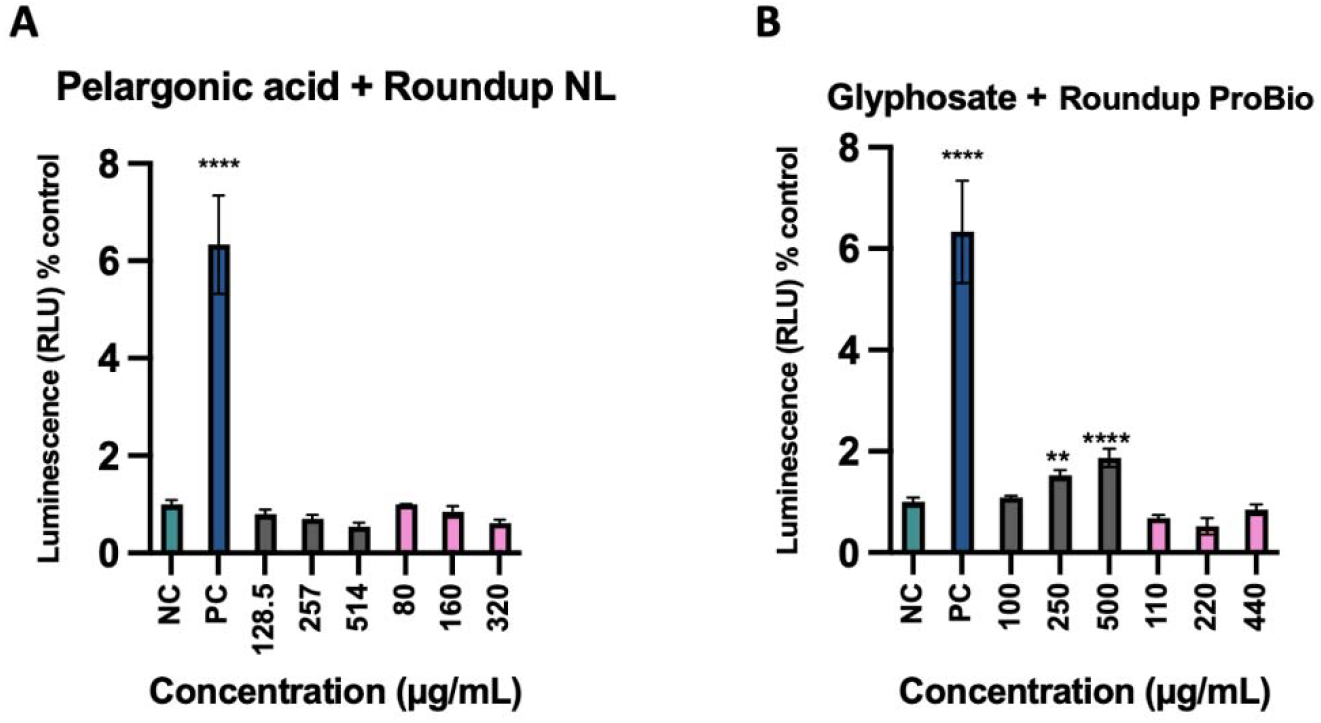
Assessing an oxidative stress response in human hepatoma HepG2 cells following treatment with pelargonic acid, Roundup NL, glyphosate and Roundup ProBio. Levels of H_2_O_2_ production as an indicator of oxidative stress was assessed in HepG2 cells exposed to pelargonic acid (**A**; grey bars), Roundup NL (**A**; pink bars), glyphosate (**B**; grey bars) and Roundup ProBio (**B**; pink column) and compared to untreated negative control (NC; green bars) and positive control (PC, 50 µM menadione; blue bars) cultures. Results are expressed as relative light units (RLU) from the luciferase reporter ROS-Glo H_2_O_2_ detection system. The assay was performed in triplicate and data are expressed as mean ± SD (standard deviation) of 3 independent replicates. ** p <0.01, ****p <0.0001 in a Šidák test which followed a one-way ANOVA.

DNA damage was assessed using the alkaline comet assay (Singh et al., 1988). The results (Figure 4A) showed that pelargonic acid, at LC10, gave rise to a modest but significant increased the percentage of DNA in the comet tail (6.50%) relative to untreated cells (4.83%). Roundup NL, tested at an equivalent concentration, caused only a marginal and non-significant increase (5.33%). This suggests that while pelargonic acid is potentially genotoxic at sub-cytotoxic doses, its commercial formulation Roundup NL did not elicit a significant increase in DNA damage under the same conditions. No significant increase of DNA in the comet tail was observed with either glyphosate alone or the GBH Roundup ProBio at their respective LC10 (Figure 4B) showing that neither substance caused DNA damage in this cell line at the sub-cytotoxic concentration tested.

**Figure 4.**
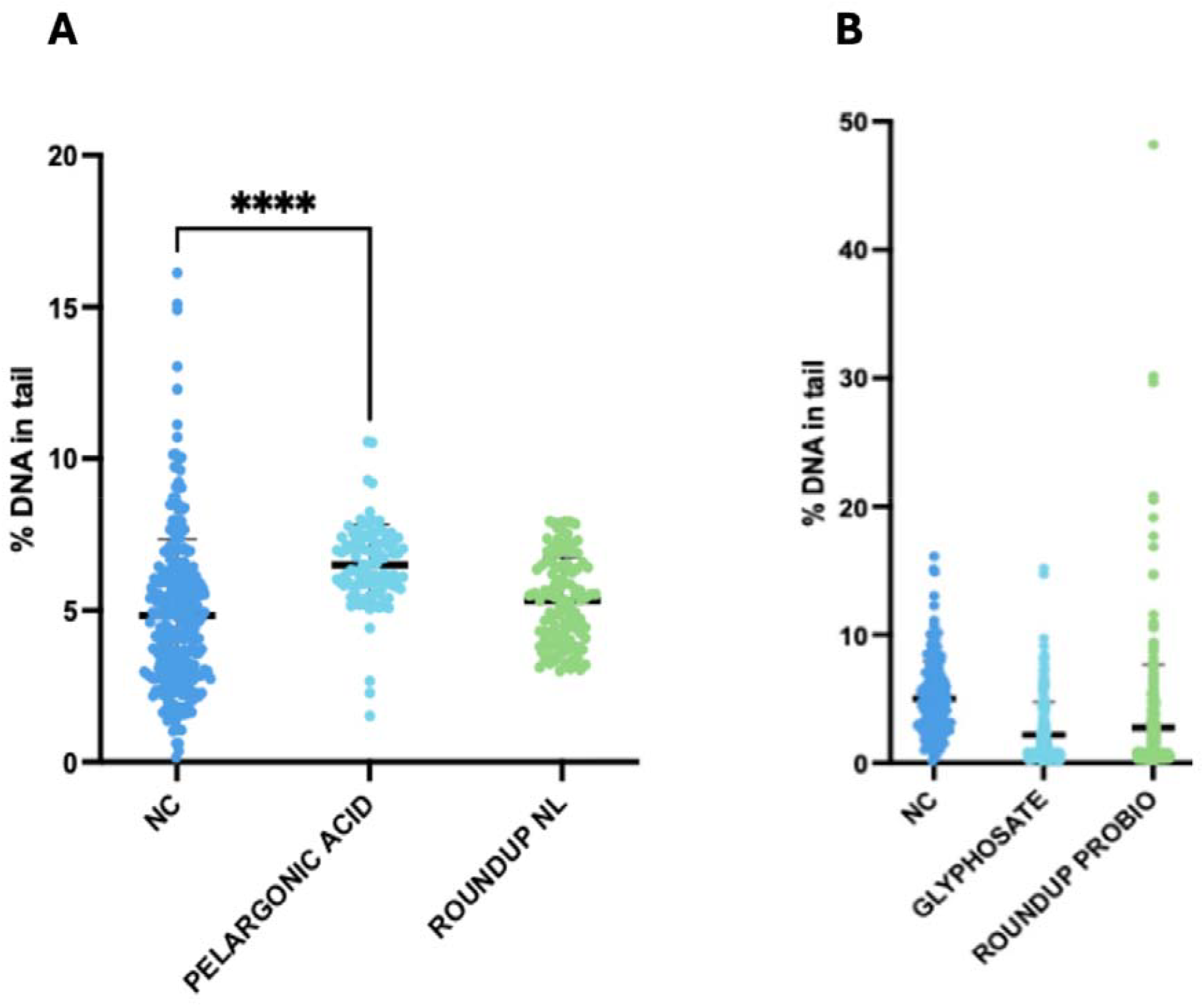
Assessment of DNA damage in HepG2 hepatoma cells treated with pelargonic acid, Roundup NL, glyphosate and Roundup ProBio using a comet assay system. Exposures were at the respective LC10 values for each test substance and compared to negative control, untreated cells (NC; dark blue). (**A**) pelargonic acid (light blue) and Roundup NL (green). (**B**) glyphosate (light blue) and Roundup ProBio (PROBIO; green). Each dot represents the comet assay result from a single cell. Exposure to pelargonic acid resulted in a significant rise in DNA damage as indicated by an increase in the percentage of DNA in the comet tail. The assay was performed in duplicate with over 100 comets scored per compound. ****p <0.0001 in a Šídák multiple comparison test which followed a one-way ANOVA.

As Roundup NL was found to be the most cytotoxic of the substances tested in HepG2 cells, the genotoxicity and cellular stress induction potential of this formulation was further evaluated in the ToxTracker assay system (Figure 5). This commercial glyphosate-free formulation did not activate the genotoxic markers Bscl2-GFP or Rtkn-GFP beyond the 2-fold threshold, indicating no direct DNA damage. A weak Rtkn-GFP signal was observed in the absence of metabolic activation but remained below the genotoxicity cut-off. However, Roundup NL did activate reporters associated with oxidative stress and proteotoxicity. The oxidative stress marker Srxn1-GFP reached the activation threshold only without metabolic activation, and Blvrb-GFP was weakly induced in both conditions. The strongest response was observed with the Ddit3-GFP reporter, indicating consistent activation of the unfolded protein response in both the presence and absence of metabolic activation. These results suggest that Roundup NL induces cellular stress responses without directly causing genotoxicity.

**Figure 5.**
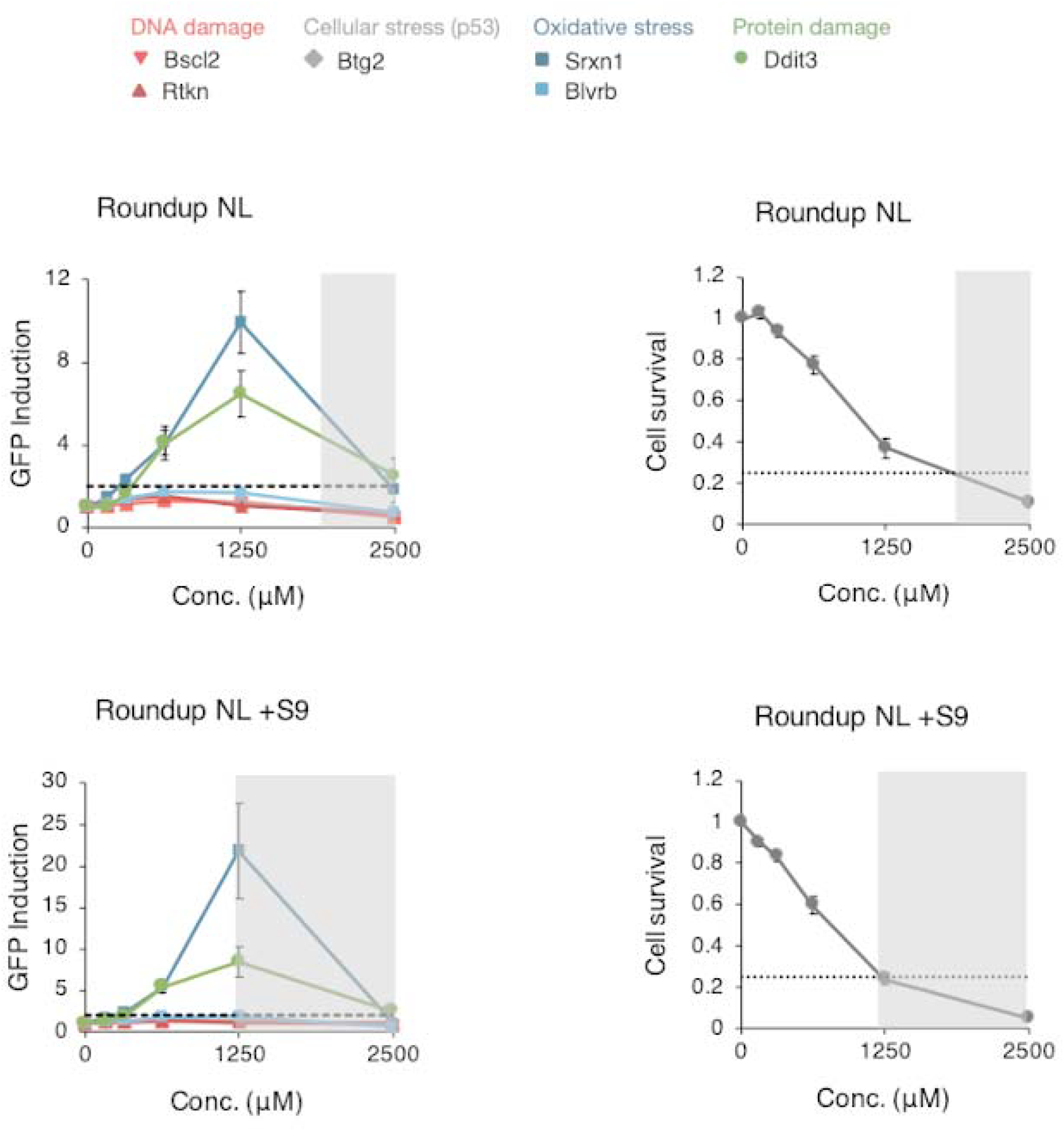
ToxTracker assay results for Roundup NL. Graphs show GFP reporter induction (left panels) and cell survival (right panels) in the ToxTracker mouse embryonic stem cell reporter assay system exposed to Roundup NL at increasing concentrations (0–2500 µM) for 24 hours, in the absence (top row) and presence (bottom row) of liver extract metabolic activation (S9). Reporter lines include Bscl2-GFP (DNA damage), Rtkn-GFP (DNA double-strand breaks), Btg2-GFP (p53 activation), Srxn1-GFP and Blvrb-GFP (oxidative stress), and Ddit3-GFP (protein damage/unfolded protein response). A transient induction of oxidative and protein stress markers (Srxn1, Ddit3) was observed below the cytotoxicity threshold (<75% cell death, grey shaded area), particularly without S9. No reporters exceeded the 2-fold activation threshold required for classification as genotoxic. Error bars indicate standard deviation from three independent experiments.

## 4. Discussion

In response to growing health and environmental concerns over GBHs, glyphosate-free alternatives such as Roundup NL, containing pelargonic acid as the active herbicidal ingredient, have been marketed as safer and more environmentally friendly. This premise is based mostly on pelargonic acid being naturally found in plants and is rapidly biodegradable rather than direct toxicological measures. In addition, toxicity assessments of commercial pelargonic acid formulations have not been determined. Although a previous investigation focused on the composition of the co-formulants present in several commercial pelargonic acid-based products (Seralini and Jungers, 2020), the study presented here is the first to assess the toxicological profile of a representative pelargonic acid formulation in established tissue culture cell model systems. Our findings challenge the safety claims that are assumed for commercial pelargonic acid formulations.

We found that pelargonic acid significantly reduced cell viability (Figure 1) and induced membrane disruption (Figure 2) in human hepatoma HepG2 cells at concentrations where glyphosate had no effect. Roundup NL, a representative commercial pelargonic acid formulation, was even more cytotoxic than pelargonic acid alone and more toxic than the GBH Roundup ProBio (Figure 1). As pelargonic acid is a nine-carbon chain lipophilic compound, it is perhaps not surprising that it can interact with and disrupt the cell plasma membrane lipid bilayer, which mimics its herbicidal mechanism of action (Preisler et al., 2026). However, the marked higher HepG2 cytotoxicity and membrane damaging capability of Roundup NL suggests that co-formulants present in this commercial formulation contribute substantially to its overall toxicity.

Unlike glyphosate, pelargonic acid and Roundup NL did not induce oxidative stress in HepG2 cells as measured by H_2_ O_2_ production (Figure 3). Glyphosate was the only compound in this study that significantly elevated ROS levels in HepG2 cells, highlighting different toxicological profiles. Nevertheless, pelargonic acid and its formulation Roundup NL were more damaging to cell membranes, indicating distinct mechanisms of cytotoxicity.

Alkaline comet assays in HepG2 cells revealed that pelargonic acid caused a modest but significant increase in DNA damage at sub-cytotoxic concentrations, while Roundup NL did not, suggesting that the formulation may modulate or mask genotoxic effects of the active ingredient (Figure 4). This contrasts with EFSA’s 2021 conclusion that pelargonic acid is unlikely to be genotoxic (EFSA, 2025). Evidently, more research is required to assess the genotoxic potential of pelargonic acid and its commercial formulations in different cell types.

The ToxTracker assay system based on mouse embryonic stem cells, confirmed that Roundup NL is not directly genotoxic, but triggered responses linked to oxidative stress and protein damage, particularly increased expression of the Ddit3-GFP genetic marker, indicating ER stress or impaired proteostasis (Figure 5). This outcome is similar to that seen with the GBH Roundup ProBio, which also elicited an oxidative stress and unfolded protein damage response in the ToxTracker assay (Mesnage et al., 2022b). Oxidative stress induced by Roundup NL in the ToxTracker assay system, contracts with a lack of such a response in HepG2 cells (Figure 3). This highlights that oxidative stress response pathways can be cell type and assay dependent, requiring the need for assessment in multiple cell types and systems. Nevertheless, the sub-lethal effects observed with Roundup NL we observed in the ToxTracker assay may have biological relevance, especially under chronic or repeated exposure scenarios.

The higher toxicity of Roundup NL relative to its declared active ingredient underscores the importance of assessing full commercial formulations, not just isolated pesticidal active ingredients, in toxicological evaluations. This aligns with findings from a previous study, which reported the presence of heavy metals and polycyclic aromatic hydrocarbons in pelargonic acid-based herbicides (Seralini and Jungers, 2020), highlighting the role of undisclosed co-formulants in enhancing toxicity.

Our investigation has a number of limitations that need to borne in mind. First, the various toxicological measures were only conducted in HepG2 cells. These need to be repeated with other cell types as sensitivity to toxicants can vary markedly between tissues. Second, long-term toxicity studies in a regulatory accepted animal model system (e.g. rodents) need to be undertaken. This is needed to evaluate if the cytotoxicity underpinned by cell membrane and DNA damage seen in HepG2 cells with pelargonic acid and Roundup NL, and an oxidative stress and protein damage response upon exposure to Roundup NL in the ToxTracker system, lead to organ damage including carcinogenesis.

Pelargonic acid is rapidly degraded and thus perceived as having a low potential in inducing negative environmental impacts (Preisler et al., 2026). However, there are two scenarios where human exposure can potentially have significant health implications. First, those using pelargonic acid-based herbicides occupationally (e.g. farm workers, groundsman) risk higher exposure through inhalation and dermal absorption as well as via food stuffs. Second, like GBHs pelargonic acid-based herbicides are also used as pre-harvest desiccants, which will lead to foodstuffs being contaminated with not only pelargonic acid but also the formulation co-formulants, which can also be toxic.

## 5. Conclusion

Overall, our findings identify pelargonic acid and Roundup NL as potential hazards to health requiring further investigation to assess more accurately their toxicological profile under real-world exposure scenarios. Our results indicate that commercial glyphosate-free herbicide formulations based on pelargonic acid are not necessarily inherently non-toxic and in certain circumstances may be as toxic as GBHs at least under acute exposure situations and that further *in vivo* studies are needed to determine their true level of safety. Roundup NL demonstrated greater cytotoxicity than either pelargonic acid, glyphosate or Roundup ProBio GBH in HepG2 cells. Roundup NL also induced a marked cell stress-inducing effect in the ToxTracker assay system as found for Roundup ProBio (Mesnage et al., 2022b). The toxicity we observed with pelargonic acid and Roundup NL also questions the assumption that alternatives derived from natural sources are non-toxic, as has been found for other substances such as pyrethrin and nicotine insecticides (Poole and Schaffer, 2024; Daraban et al., 2023). Furthermore, our findings with Roundup NL add to the large and growing body of evidence that pesticide commercial formulations are generally more toxic than stipulated pesticidal active ingredients alone (Mesnage and Antoniou, 2018; Mesnage et al., 2019), and thus regulatory frameworks must evolve to consider full formulation short and long-term toxicity and ensure safety assessments reflective of real-world exposures.

## Author contributions

**Scarlett Ferguson:** Formal analysis, Investigation. **Robin Mesnage:** Conceptualization, Data curation, Formal analysis, Investigation, Methodology, Writing – original draft, Writing – review and editing. **Michael N Antoniou:** Conceptualization, Data curation, Formal analysis, Funding acquisition, Methodology, Writing – original draft, Writing – review and editing.

## Statements and Declarations Competing Interests

RM and MNA have served as consultants on glyphosate risk assessment issues as part of litigation in the US over glyphosate-based herbicide health effects. SF declares no competing interests.

## Acknowledgments

This work was funded by the Sustainable Food Alliance (USA) whose support is gratefully acknowledged.

